# Repurposing diphenylbutylpiperidine-class antipsychotic drugs for host-directed therapy of *Mycobacterium tuberculosis* and *Salmonella enterica* infections

**DOI:** 10.1101/2021.06.05.447191

**Authors:** MT Heemskerk, CJ Korbee, J Esselink, C Carvalho dos Santos, S van Veen, IF Gordijn, F Vrieling, KV Walburg, CG Engele, K Dijkman, L Wilson, FAW Verreck, THM Ottenhoff, MC Haks

## Abstract

The persistent increase of multidrug-resistant (MDR) *Mycobacterium tuberculosis* (*Mtb*) infections negatively impacts Tuberculosis (TB) treatment outcomes. Host-directed therapies (HDT) pose an complementing strategy, particularly since *Mtb* is highly successful in evading host-defense by manipulating host-signaling pathways. Here, we screened a library containing autophagy-modulating compounds for their ability to inhibit intracellular *Mtb*-bacteria. Several active compounds were identified, including two drugs of the diphenylbutylpiperidine-class, Fluspirilene and Pimozide, commonly used as antipsychotics. Both molecules inhibited intracellular *Mtb* in pro- as well as anti-inflammatory primary human macrophages in a host-directed manner and synergized with conventional anti-bacterials. Importantly, these inhibitory effects extended to MDR-*Mtb* strains and the unrelated intracellular pathogen, *Salmonella enterica* serovar Typhimurium (*Stm*). Mechanistically Fluspirilene and Pimozide were shown to regulate autophagy and alter the lysosomal response, partly correlating with increased bacterial localization to autophago(lyso)somes. Pimozide’s and Fluspirilene’s efficacy was inhibited by antioxidants, suggesting involvement of the oxidative-stress response in *Mtb* growth control. Furthermore, Fluspirilene and especially Pimozide counteracted *Mtb*-induced STAT5 phosphorylation, thereby reducing *Mtb* phagosome-localized CISH that promotes phagosomal acidification.

In conclusion, two approved antipsychotic drugs, Pimozide and Fluspirilene, constitute highly promising and rapidly translatable candidates for HDT against *Mtb* and *Stm* and act by modulating the autophagic/lysosomal response by multiple mechanisms.

## Introduction

Tuberculosis (TB), an infectious disease caused by the pathogen *Mycobacterium tuberculosis* (*Mtb*), generally manifests as pulmonary disease, although any other organ can be affected. Infection is transmitted by aerosol borne *Mtb* which is phagocytosed by alveolar macrophages [1]. Currently, 23% of the world’s population is estimated to be latently infected with the bacillus, from which around 5-10% will progress towards developing active disease during their lifetime [2]. In 2019, this led to around 10 million new TB cases, and an unacceptable number of over 1.4 million deaths [2]. Major efforts addressing new TB vaccine candidates are in progress [3], mainly because the only TB vaccine currently available, Bacillus Calmette-Guérin (BCG) can protect infants against severe forms of TB, but is clearly insufficient in preventing pulmonary TB in adults [4].

Another major class of human pathogens is the genus of *Salmonella*, which contains several serovars that cause significant global morbidity and mortality. Typhoid and paratyphoid fever are caused by the *Salmonella enterica* serotype Typhi and Paratyphi, respectively, while nontyphoidal salmonellosis (gastroenteritis) is related to other serovars, such as *Salmonella enterica* serovar Typhimurium (*Stm*) [5]. More than a billion people are affected annually by these infections, leading to several hundreds of thousands of deaths [5, 6].

A major difficulty in TB control is the increase in infections with MDR- and extensive-drug resistant TB (XDR-TB), with only 39% being treated successfully in case of the latter [2]. Also, the emergence of drug-resistant *Salmonella* is a major concern, urging the development of new therapeutics [7]. Although numerous antibiotics against TB are currently in clinical trials and a few new antibiotics for the treatment of MDR-TB and XDR-TB, Bedaquilin, Linezolid and Pretomanid [8, 9], have recently been approved, several reasons encourage the search for alternative treatment strategies [10]. Barriers such as toxicity, long treatment duration, high costs and social stigma are hampering treatment adherence, thus contributing to the emergence of antibiotic resistance [11]. Additionally, the inevitable emergence of future resistance against new antibiotics will negatively impact the global goal of eradicating TB, prompting the need for complementary strategies. Host-directed therapy (HDT) might prove a successful contributor towards this goal since it is expected to shorten treatment duration, has the potential to synergize with anti-bacterial therapy, could eradicate dormant bacteria which often escape conventional antibiotic treatment, and should be equally effective against drug-susceptible and drug-resistant *Mtb* [12]. An additional advantage is that bacterial resistance against HDT is unlikely to be a problem because single bacterial mutations will be insufficient to counteract multiple host mechanisms simultaneously [13]. As a first step towards HDT, drug repurposing is an attractive strategy because it could rapidly create a pipeline for new treatment modalities [14].

The reason for *Mtb*’s persistence in human host cells is thought to primarily be due to its ability to modify host signaling and effector pathways to its own advantage. Upon the first contact with phagocytic cells, host cell manipulation is initiated by using extracellular receptors, for example the induction of anti-inflammatory IL-10 via TLR2 and the reduction of pro-inflammatory cytokine production via a pathway initiated by the binding of mycobacterial mannose-capped lipoarabinomannan to the mannose receptor [15, 16]. Following phagocytosis, *Mtb* halts the process of phagosome maturation by preventing fusion with lysosomes [17]. Additionally, the bacterium induces the depletion of the vacuolar ATPase (v-ATPase) from the *Mtb* phagosome, which results in decreased lysosomal acidification and bacterial degradation and therefore decreased antigen processing and antigen presentation, impairing both innate and adaptive immune responses [18]. Furthermore, the mycobacterial secretion machinery, such as the ESAT-6 secretion system-1 (ESX-1) encoded by the region of difference 1 (RD1) that is present in virulent *Mtb* strains but absent from BCG, is of vital importance in manipulating host defense mechanisms targeting the bacterium, exemplified by the inhibition of autophagy by ESX-1 secretion-associated protein B (EspB) [19, 20]. *Salmonellae* also possess numerous mechanisms through which it manipulates infected host cells, including the activation of the host kinase Akt1 to inhibit phagosomal maturation via the Akt1/AS160 pathway [21]. Thus, numerous bacterial effector molecules, including constituents of the bacterial cell wall, can impair the host cell response and promote the bacteria’s survival.

Various studies, including our own, have aimed to identify druggable host target molecules for HDT [22–25]. Many of these studies underscored the importance of autophagy in controlling the intracellular *Mtb* lifecycle [25–27]. Autophagy regulates cellular homeostasis through recycling intracellular cargo, including protein aggregates, damaged organelles and intracellular pathogens, and delivering these to lysosomes for destruction [28]. Autophagy selectively targeting intracellular pathogens, also termed xenophagy, has been shown to restrict replication and survival of both *Mtb* and *Salmonellae* [29], and is activated by various host molecules, such as IFN-γ [30], TNF-α [31], and extracellular DNA sensor stimulator of interferon genes (STING) [32], for which we showed the DNA damage-regulated autophagy modulator 1 (DRAM1) to play an important role [27]. Several small molecule activators of autophagy have been reported to lower intracellular *Mtb* burden, including metformin, statins, nitazoxanide, gefitinib and imatinib [14].

Although metformin treatment of patients suffering from diabetes mellitus (DM) lowered the prevalence of latent tuberculosis infection (LTBI) [33], it failed to show an anti-TB effect in DM patients with pulmonary active TB [34, 35]. Statins have similarly been shown to lower LTBI prevalence [33], but in addition have pleiotropic effects on the host response, which can both positively and negatively impact treatment outcome [36]. Nitazoxanide has been shown to have host-directed activity against *Mtb* by inducing autophagy, although it also has direct antimicrobial activity [37]. Tyrosine kinases, including those sensitive to Imatinib such as ABL1, have been found to play a role in *Mtb* infection by several groups including ours [22, 38, 39]. Corroborating these findings, Imatinib has been shown to display anti-mycobacterial activity *in vivo* in mice [40] and is currently being evaluated in a phase 2 trial.

Drugs with efficacy towards controlling intracellular infections with low toxicity are still in short supply. Here, we screened a library of compounds with defined autophagy-inducing or -inhibitory activity using our previously described, human cell based (*Mtb*-MelJuSo) intracellular infection model [22]. Several promising candidates were identified, among which two antipsychotic drugs of the diphenylbutylpiperidine-class [41], Fluspirilene and Pimozide, were found as particularly interesting hits.

We found that Fluspirilene and Pimozide employ multiple mechanisms converging mostly on the autophagosomal/lysosomal response pathway to control different species of intracellular bacteria. Pimozide was particularly able to induce reactive oxygen species (ROS) which are of vital importance in the host defense response. Furthermore, Fluspirilene and particularly Pimozide inhibited STAT5-dependent *Mtb*-induced CISH-mediated degradation of phagosomal v-ATPase, underscoring the feasibility of targeting this recently uncovered mechanism. In conclusion we have uncovered potent HDT activity of two antipsychotics against (drug resistant) intracellular *Mtb* and *Stm* in human cells.

## Materials and methods

### Chemicals

The Screen-Well Autophagy Library version 1.2 (http://www.enzolifesciences.com/BML-2837/screen-well-autophagy-library/) was purchased from Enzo Life Sciences, Brussels, Belgium. H-89 diHCl (H-89), Fluspirilene, Pimozide, Ebselen, NG-Methyl-L-arginine acetate salt (L-NMMA), MitoTEMPO, Phorbol 12-myristate 13-acetate (PMA), N-acetyl cysteine (NAC), tert-butylhydroperoxide (TBHP) and Rifampicin were purchased from Sigma-Aldrich, Zwijndrecht, The Netherlands. Torin-1 and Isoniazid were purchased from SelleckChem, Munich, Germany. Hygromycin B was acquired from Life Technologies-Invitrogen, Bleiswijk, The Netherlands and Gentamicin sulfate was bought from Lonza BioWhittaker, Basel, Switzerland.

### Antibodies

Rabbit anti-phospho-STAT5 (RRID:AB_823649), rabbit anti-CISH (RRID:AB_11178524) and rabbit polyclonal anti-TFEB (RRID:AB_11220225) were all bought from Cell Signaling Technology, Leiden, The Netherlands. Phalloidin-iFluor 647 and Phalloidin-iFluor 405 were obtained from Abcam, Cambridge, United Kingdom. Goat anti-rabbit IgG (H+L) AlexaFluor647 conjugate (RRID:AB_2536101) was purchased from ThermoFisher Scientific, Breda, The Netherlands.. Anti-human CD11b-PE (RRID:AB_395789) and anti-human CD1a-BV605 (RRID:AB_2741933) were acquired from BD BioSciences, Vianen, The Netherlands and anti-human CD14-FITC (RRID:AB_830677) and anti-human CD163-AF647 (RRID:AB_2563475) from BioLegend, San Diego, CA, USA.

### Cell culture

HeLa and MelJuSo cell lines were maintained at 37°C/5% CO_2_ in Gibco Iscove’s Modified Dulbecco’s Medium (IMDM, Life Technologies-Invitrogen) supplemented with 10% fetal bovine serum (FBS; Greiner Bio-One, Alphen a/d Rijn, The Netherlands), 100 units/ml Penicillin and 100 µg/ml Streptomycin (both Life Technologies-Invitrogen) (complete IMDM) as described previously [22, 23].

Buffy coats were obtained from healthy donors after written informed consent (Sanquin Blood Bank, Amsterdam, The Netherlands). Peripheral blood mononuclear cells (PBMCs) were purified using density gradient centrifugation over Ficoll-Paque and monocytes isolated with subsequent CD14 MACS sorting (Miltenyi Biotec, Bergisch Gladsbach, Germany) as described previously [22, 23]. Monocytes were differentiated into pro-inflammatory (Mφ1) or anti-inflammatory (Mφ2) macrophages with 5 ng/ml of granulocyte-macrophage colony-stimulating factor (GM-CSF; Life Technologies-Invitrogen) or 50 ng/ml macrophage colony-stimulating factor (M-CSF; R&D Systems, Abingdon, UK), respectively, for 6 days with a cytokine boost at 3 days, as previously reported [42]. Cells were cultured at 37°C/5% CO_2_ in Gibco Roswell Park Memorial Institute (RPMI) 1640 medium or RPMI 1640 (Dutch modified) (Life Technologies-Invitrogen) supplemented with 10% FBS and 2 mM L-alanyl-L-glutamine (GlutaMAX) (PAA, Linz, Austria), 100 U/ml penicillin and 100 µg/ml streptomycin (complete RPMI) at a density of 1×10^6^ cells per ml in T75 flasks (Sigma-Aldrich). Macrophages were harvested using Trypsin-EDTA 0.05% (ThermoFisher Scientific) and scraping and macrophage differentiation was evaluated by cell surface marker expression of CD11b, CD1a, CD14 and CD163 implementing flow cytometry and by quantification of IL-10 and IL-12p40 secretion using ELISA following 24-hour stimulation in the presence or absence of 100 ng/ml of lipopolysaccharide (LPS; InvivoGen, San Diego, United States).

Bronchoalveolar lavage (BAL) samples from purpose-bred, Indian-type rhesus macaques in the present study, were used as they were available occasionally, when an animal happened to be indicated for ketamine-sedation and euthanasia for veterinary and animal welfare reason. Thus, the availability of sample was exploited as it occurred, beyond any legal requirement for prior approval of protocol as there was no pre-existing study plan nor any discomfort afflicted to animals for the sake of a research objective. Using a bronchoscope for instillation and recovery, cells were harvested by flushing with 3 consecutive volumes of 20 mL of sterile, pre-warmed, isotonic saline solution. Samples were immediately put on ice and kept cold until further processing. To isolate the cellular fraction, the BAL was filtered over a 100 μm filter and spun down at 400g for 10 min at 4°C. To enrich BAL cells for alveolar macrophages (AMφ), BAL cells were resuspended in complete RPMI and incubated for 4 hours at 37°C/5% CO_2_ in a T75 flask after which non-adherent cells were discarded. Adherent cells were washed with PBS and harvested using Trypsin-EDTA 0.05% and scraping. Cells were counted using Türk solution, spun down by centrifugation at 400g for 10 min, resuspended in complete RPMI and seeded at 300,000 per ml for downstream application. NHP samples were obtained at the Biomedical Primate Research Centre (BPRC), Rijswijk, the Netherlands. BPRC is licensed by the Dutch authority to breed non-human primates and to use them for research in areas of life-threatening and disabling diseases without suitable alternatives. BPRC complies to all relevant legislation with regard to the use of animals in research; the Dutch ‘Wet op de Dierproeven’ and the European guideline 2010/63/EU. BPRC is AAALAC accredited since 2012.

### Bacterial infection of cells

*Mtb* (wild-type H37Rv or DsRed-expressing H37Rv) [22] was cultured in Difco Middlebrook 7H9 broth (Becton Dickinson, Breda, the Netherlands) supplemented with 10% ADC (Becton Dickinson), 0.05% Tween 80 (Sigma-Aldrich), and with the addition of 50 µg/ml Hygromycin B in the case of DsRed-expressing H37Rv. *Mtb* suspensions were prepared from a running *Mtb* culture, which was one day prior to infection diluted to a density corresponding with early log-phase growth (OD600 of 0.25). *Stm* strain SL1344 was cultured in Difco lysogeny broth (LB) (Becton Dickinson). *Stm* was grown overnight in LB, subsequently diluted 1:33 in fresh LB and used after approximately 3 hours of incubation, when log-phase growth was achieved (OD600 of 0.5). Bacteria (or liquid broth at equal v/v for mock-infection) were diluted in complete IMDM or complete RPMI without antibiotics for infecting cell lines or primary cells, respectively. We consistently used a multiplicity of infection (MOI) of 10 for both strains. Cell lines and primary cells, which had been seeded at a density of 10,000 or 30,000 cells per well, respectively, in 96-well flat-bottom plates 1 day prior to infection, were inoculated with 100 μl of the bacterial suspension. Cells were subsequently centrifuged for 3 min at 800 rpm and incubated at 37°C/5% CO_2_ for 20 min in case of *Stm* infection or 60 min in case of *Mtb* infection. Extracellular bacteria were then washed away with culture medium containing 30 μg/ml Gentamicin sulfate, incubated for 10 min at 37°C/5% CO2, followed by replacement with medium containing 5 μg/ml Gentamicin sulfate and, if indicated, chemical compounds until readout. MOI of the bacterial inoculum was confirmed by a standard colony-forming unit (CFU) assay.

### Chemical compound treatment

Cells were treated for 24 hours, unless indicated otherwise, with chemical compound at a concentration of 10 µM, unless indicated otherwise, or DMSO at equal v/v in medium containing 5 μg/ml Gentamicin sulfate. Treatment regimens were designed not to exceed DMSO solvent end concentrations of 0.2%.

### Colony-forming unit (CFU) assay

Cells were lysed in H_2_O containing 0.05% SDS (ThermoFisher Scientific). Lysates of *Mtb-*infected cells were serially diluted in steps of 5-fold in 7H9 broth and 10 μl droplets were spotted onto square Middlebrook 7H10 agar plates. Plates were incubated at 37°C for 12 -14 days and bacterial colonies quantified using a microscope with a magnification of 2.5 times to enhance early detection of bacterial growth. Lysates of *Stm*-infected cells were serially diluted in LB broth, thereafter 10 μl droplets were spotted onto square LB agar plates and incubated overnight at 37°C.

### Liquid bacterial growth inhibition assay

*Stm* or *Mtb* cultures in logarithmic growth phase were diluted to an OD_600_ of 0.1 in LB broth or 7H9 broth respectively, of which 200 μl per flat-bottom 96-well was incubated with chemical compound, antibiotic or DMSO at equal v/v at indicated concentrations. *Stm* growth was measured after overnight incubation at 37°C, while *Mtb* growth was evaluated for 10 days of incubation at 37°C. Absorbance was measured by optical density at 550 nm on a Mithras LB 940 plate reader (Berthold Technologies, Bad Wildbad, Germany) while shaking between measurements.

### Flow cytometry

Infected cells were at experimental endpoint washed with 100 µl of PBS and detached by incubation in 50 µl of Trypsin-EDTA 0.05% for five minutes. Single cell suspensions were fixed by adding 100 µl of 1.5% paraformaldehyde and incubated for 60 min at 4°C. Acquisition was performed using a BD FACSLyric™ Flow Cytometer equipped with BD FACSuite software (BD Biosciences). Data was analyzed using FlowJo software v10.

### Cell viability assay

Cells seeded at a density of 30,000 cells/well in 96-well flat-bottom plates were stained in 50 μl RPMI without phenol red (Life Technologies-Invitrogen) containing propidium iodide (PI) (1:500, Sigma-Aldrich) and 100 μg/ml of Hoechst 33342 (Sigma-Aldrich). Cells were incubated at room temperature (RT) for 5 min. Per well 3 images were taken using a Leica AF6000 LC fluorescence microscope combined with a 20x dry objective. The number of dead cells (positive for PI) versus total number of adherent cells (positive for Hoechst) was quantified using CellProlifer version 3.0.0 [55].

### Immunostaining

Cells were seeded on poly-d-lysine coated glass-bottom (no. 1.5) 96-well plates (MatTek, Ashland, Massachusetts, United States), pre-washed with PBS, at a density of 30,000 per well. After overnight incubation, cells were infected with DsRed-expressing *Mtb* at a MOI of 10 as previously described. At the indicated experimental endpoint, cells were washed three times with PBS and fixed for 60 min at RT using 1% methanol-free EM-grade formaldehyde (ThermoFisher Scientific) diluted in PBS. Cells were washed with PBS and remaining reactive formaldehyde was quenched using 100 μl of Glycine solution (1.5 mg/ml in PBS) for 10 min at RT. Cells were permeabilized in 0.1% Triton-X (Sigma-Aldrich) diluted in PBS for 10 min at RT and Fc-receptors were subsequently blocked using 5% human serum (HS; Sanquin Blood bank, Amsterdam, The Netherlands) for 45 min at RT. After removal of the 5% HS, cells were stained with primary antibody diluted in 5% HS for 30 min at RT, washed three times with 5% HS and incubated with secondary antibody in 5% HS for 30 min at RT in the dark. After washing three times with 5% HS, cells were counterstained with 50 ul of 2 μg/ml Hoechst 33342 and Phalloidin for 30 minutes at RT in the dark. Images, at least 3 per well, were acquired using a Leica TCS SP8 X WLL confocal system and 63X oil immersion objective. Hybrid detectors were used with a time gate to switch off data collection during the pulse. The fluorescent dyes LysoTracker Deep Red (ThermoFisher Scientific) (75 nM) and CYTO-ID 2.0 (Enzo LifeSciences) (1:500) were added to the cells 30 minutes prior to treatment endpoint. Cells were fixed at the experimental endpoint and counterstained with Hoechst and Phalloidin as described above except for permeabilization.

Colocalization analysis was performed as follows. Image background was subtracted using the rolling ball (20-pixel radius) algorithm with Fiji software [43]. CellProlifer 3.0.0 was used to first correct for non-homogenous illumination if necessary, then for the segmentation of both the fluorescent bacteria and marker of interest using global thresholding with intensity-based de-clumping [44]. For every experiment, segmentation was performed independently with both a range of thresholds and adaptive three-class Otsu thresholding to confirm segmentation results. Then per image the overlap of *Mtb* with marker of interest was calculated as percentage of object overlap (colocalization) or the integrated intensity of the marker of interest per single nucleus or bacterium was determined. Data in Figures 3C and 4C are shown for 4 individuals, while two donors could not be analyzed due to high background levels.

### Reactive oxygen species (ROS) / Reactive nitrogen species (RNS) assay

Cells were seeded in complete RPMI in Corning 96-well Special Optics Flat Clear Bottom Black Polystyrene TC-Treated Microplates (Sigma-Aldrich) at a density of 40,000 per well. After overnight incubation at 37°C/5% CO_2_, cells were washed once with PBS and incubated with 5 μM CM-H2DCFDA (Life Technologies-Invitrogen), a general oxidative stress indicator, for 30 min at 37°C/5% CO_2_ in phenol red free RPMI. Cells were washed twice with PBS followed by treatment with chemical compounds (10 µM) in 100 μl of complete RPMI without phenol red. 50 µM of TBHP was used as technical assay control. Oxidation of the probe yields a fluorescent adduct which presence was quantified every 5 min for 2 hours by measuring fluorescence using the SpectraMax i3x at 37°C. Specific settings used were 10 flashes per read with an 493/9 excitation and a 522/15 emission filter pair and a high PMT gain.

### Total RNA isolation and cDNA synthesis

Total RNA isolation was performed using TRIzol Reagent (Life Technologies-Invitrogen) according to the manufacturer’s instructions and described previously [23]. RNA yield was quantified using a DeNovix DS-11 Spectrophotometer (ThermoFisher Scientific). Total RNA (0.5 μg) was reverse transcribed using SuperScript IV Reverse Transcriptase (Life Technologies-Invitrogen). Briefly, RNA samples were first incubated at 65°C for 5 min in the presence of 0.5 mM dNTPs and 2.5 µM oligo(dT)20 (Life Technologies-Invitrogen). Subsequently, cDNA synthesis was initiated by adding a master mix containing 1x first strand buffer, 5 mM DTT, 40 U RNaseOUT (ThermoFisher Scientific) and 200 U SuperScript IV and incubating at 50-55°C for 10 min followed by inactivation of the reverse transcriptase at 80°C for 10 min.

### TaqMan qPCR

Multiplex quantitative polymerase chain reaction (qPCR) was carried out using a QuantStudio 6 Flex Real-Time PCR System (ThermoFisher Scientific) as described previously [23]. qPCR reactions were performed in a final volume of 25 µl containing 1x TaqMan Universal PCR Master Mix, No AmpErase UNG, 0.5x CISH-FAM TaqMan primers (Hs00367082_g1, ThermoFisher Scientific), 0.5x GAPDH-VIC TaqMan primers ((Hs02758991_g1, ThermoFisher Scientific), and 5 µl cDNA. Thermal cycling conditions were 1 cycle of 2 min/50°C and 10 min/95°C, followed by 40 cycles of 15 s/95°C and 1 min/60°C. The threshold cycle (Ct) values of CISH transcripts were normalized to GAPDH by the 2^-ΔΔCT^ algorithm method [45]. Relative expression levels were calculated by applying the formula ((2^-ΔCT(Target gene)^)/(2-^ΔCT(GAPDH)^)).

### Statistics

Statistics were performed as described previously [23]. Normal distribution of the data was tested using the Shapiro-Wilk test. If the data was normally distributed, unpaired t-test with Welch’s correction or one-way ANOVA with Dunnett’s multiple comparison test were applied when assessing differences between 2 and 3 or more groups of unpaired data, respectively. Normally distributed paired samples were analyzed using repeated measure (RM) one-way ANOVA with Dunnett’s multiple comparison test and Geisser Greenhouse correction. Two-way ANOVA with Tukey’s multiple comparison test was used when the effect of two independent variables was tested simultaneously. Assessing differences between 2 or more groups of unpaired observations from not normally distributed data was performed using Kruskal-Wallis test, while paired observations were tested using the Wilcoxon matched-pairs signed rank test with post-hoc Benjamini-Hochberg correction. Analyses were performed using GraphPad Prism 8.

### Data availability

The datasets generated and analyzed during the current study are not deposited in external repositories but are included as supplementary file or available from the corresponding author on reasonable request.

## Results

### Compounds were identified that contain autophagy-regulating activity and repurposing potential for HDT against *Mtb* and *Stm*

By employing our previously described flow-cytometry-based assay using *Mtb*-infected human MelJuSo cells [22], we screened the Screen-Well Autophagy library that includes clinically approved molecules, to find new drugs with HDT activity against intracellular *Mtb. Mtb*-infected human cells were treated during 24h with compounds. The PKA/PKB-Akt1 inhibitor H-89 was included as a positive control based on our earlier work [21, 22]. We identified 25 compounds that strongly reduced bacterial burden (<35% infected cells compared to control DMSO). From these 25 molecules we discontinued 15 because of undesirable significant host cell toxicity (cell-yield <85% after 24h treatment compared to control DMSO), and 2 because of a current lack of clinical approval (SB-216763 and Licochalcone A). Two additional compounds were not pursued further since their *in vivo* adjunct therapy potential had already been shown previously, independently validating our discovery approach (Chloroquine and its metabolite Hydroxychloroquine) [46] (**Figure 1A, and Table S1**).

**Figure 1.**
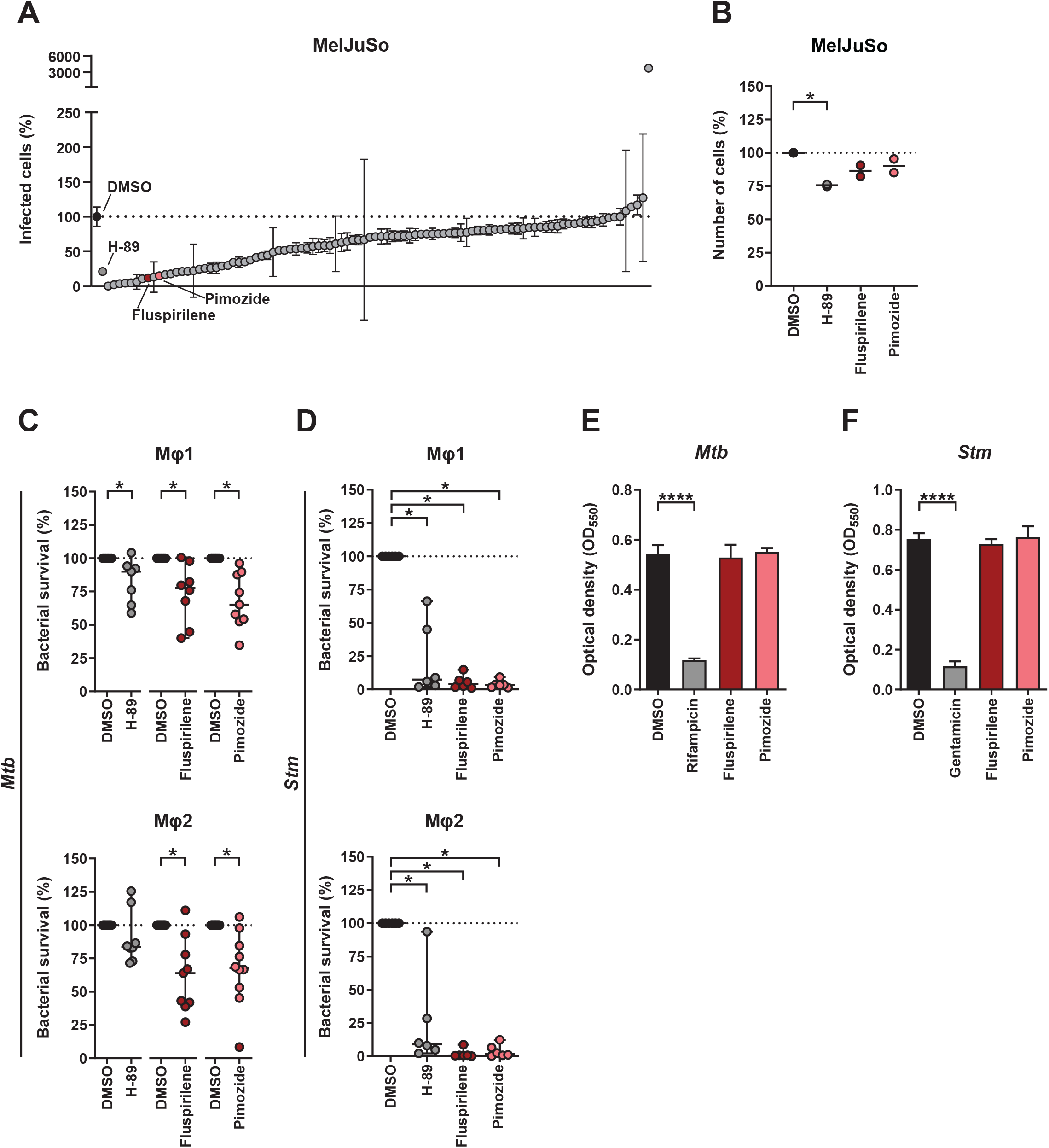
Identification of compounds with autophagy-regulating activity and repurposing potential for HDT against *Mtb* and *Stm*. **A.** MelJuSo-*Mtb* model-based screening results of the Screen-Well Autophagy library at 10 µM concentration after 24 hours of treatment sorted by effect size on the percentage of infected cells (DsRed-*Mtb* positive cells) compared to DMSO. Data points display the mean ± standard deviation of 3 replicates. The positive control H-89 and two structural analogues Fluspirilene and Pimozide are annotated. **B.** MelJuSo-*Mtb* model-based screening results of H-89, Fluspirilene and Pimozide at 10 µM concentration after 24 hours of treatment on the percentage of cells compared to DMSO. Data points display the mean of 3 replicates and represent two independent experiments. Dotted lines indicate DMSO set at 100% and median is shown for each condition. Statistical significance was tested using a RM one-way ANOVA with Dunnett’s multiple comparison test (* = p-value <0.05). **C.** CFU assay of Mφ1 (upper panel) and Mφ2 (lower panel) infected with *Mtb* and treated with 10 µM of Fluspirilene, Pimozide, H-89 as positive control or DMSO at equal v/v for 24 hours. Effect of compound treatments are shown separate since donors tested were not always identical. Each dot represents a single donor (7 and 8 for H-89, 8 and 9 for Fluspirilene and 9 and 10 donors for Pimozide were tested in Mφ1 and Mφ2, respectively) and depicts the mean of 3 or 4 replicates. Dotted lines indicate DMSO set at 100% and median + 95% confidence intervals are shown for every condition. Statistical significance was tested using Wilcoxon matched-pairs signed rank test with post-hoc Benjamini-Hochberg correction (right panel) (* = q-value <0.1). **D.** CFU assay of Mφ1 (upper panel) and Mφ2 (lower panel) infected with *Stm* and treated with 10 µM of Fluspirilene, Pimozide, H-89 as positive control or DMSO at equal v/v for 24 hours. Each dot represents a single donor (6 donors in total) and depicts the mean of 3 or 4 replicates. Dotted lines indicate DMSO set at 100% and median + 95% confidence intervals are shown for every condition. Statistical significance was tested using Wilcoxon matched-pairs signed rank test with post-hoc Benjamini-Hochberg correction (right panel) (* = q-value <0.1). **E.** *Mtb* growth in liquid culture during treatment with 10 μM of Fluspirilene, Pimozide or DMSO at equal v/v at assay endpoint, day 10. Rifampicin (20 μg/ml) was used as positive control for *Mtb* growth inhibition. Bars depict mean ± standard deviation of 3 replicates. Experiment shown is a representative of 4 independent experiments. Statistical significance of treatment versus DMSO was tested using a one-way ANOVA with Dunnett’s multiple comparisons test (**** = p-value <0.0001). **F.** *Stm* growth in liquid culture during treatment with 10 μM of Fluspirilene, Pimozide or DMSO at equal v/v. Gentamicin (50 μg/ml) was used as positive control for *Stm* growth inhibition. Bars depict mean ± standard deviation of 3 replicates. Experiment shown is a representative of 2 independent experiments. Statistical significance of treatment versus DMSO was tested using a one-way ANOVA with Dunnett’s multiple comparisons test (**** = p-value <0.0001).

In our effort to focus on new candidates it was of interest that 2 of the 6 remaining compounds, notably Fluspirilene and Pimozide, are structural analogues of the diphenylbutylpiperidine-class of antipsychotic drugs. Both drugs have an extensive clinical safety profile and are already for several decades used to treat multiple disorders. To explore and compare mechanistics of the analogues, we decided to focus on these two drugs only in this study, while detailed work on the other hit compounds will be described elsewhere.

We observed no cellular toxicity following treatment with Fluspirilene and Pimozide, while H-89 treatment affected cell numbers slightly (**Figure 1B**). To validate these initial screening results further in a physiologically more relevant *in vitro* model, we generated primary GM-CSF-derived pro-inflammatory macrophages (Mφ1) and M-CSF-derived anti-inflammatory macrophages (Mφ2) [22, 42, 47], and tested the potential of the two diphenylbutylpiperidines to inhibit outgrowth of intracellular *Mtb* (using a classical colony forming unit (CFU) assay as read-out (**Figure 1C**). Both Fluspirilene and Pimozide induced a significant decrease of *Mtb* outgrowth in Mφ1 and particularly Mφ2 (median reduction of bacterial outgrowth of 22% and 41% for Fluspirilene and 35% and 73% for Pimozide, respectively) (**Figure 1C**).

To investigate potential broader-range applicability of Fluspirilene and Pimozide, we investigated their intracellular bacterial growth inhibitory activity against a different class of intracellular bacteria, using *Stm*-infected human Mφ (**Figure 1D**). H-89 was again included as a positive control (it displays greater activity against *Stm* than *Mtb*) [21, 22]. Interestingly, both drugs vastly reduced *Stm* outgrowth (median reduction of bacterial outgrowth > 96% and > 98% in Mφ1 and Mφ2, respectively), suggesting these HDT drugs could be more broadly applicable.

To exclude that Fluspirilene and Pimozide acted in a direct anti-bacterial manner, extracellular *Stm* and *Mtb* were treated with Fluspirilene and Pimozide in liquid broth, in equal concentrations (10 µM) as used in the cell-based infection models. Fluspirilene and Pimozide did not affect *Mtb* or *Stm* growth, whereas control antibiotics Rifampicin (*Mtb*) and Gentamicin (*Stm*) inhibited bacterial growth (**Figures 1E and F**).

Taken together, we identified two clinically approved structural analogues displaying reported autophagy-inducing capacities as novel safe candidate molecules for HDT, that inhibited intracellular *Mtb* as well as *Stm* in both pro- and anti-inflammatory human primary macrophages.

### Fluspirilene and Pimozide HDT activity against intracellular *Mtb* was confirmed cross-species of host and mycobacterium

Since HDT drugs act on host rather than bacterial targets, they should inhibit outgrowth of drug-susceptible and multi-drug resistant (MDR)-*Mtb* strains similarly. To verify this, we measured the effect of the two diphenylbutylpiperidines on CFU outgrowth in human Mφ2 (we focused on Mφ2 since we found similar efficacy in both Mφ1 and Mφ2). The cells were infected with either *Mtb* Dutch outbreak strain 2003-1128 or *Mtb* Beijing strain 16319, both MDR-*Mtb*-strains. Indeed, treatment with Fluspirilene and Pimozide significantly inhibited bacterial outgrowth of both MDR-*Mtb* strains highly efficiently (**Figure 2A**), thereby emphasizing their clinical relevance.

**Figure 2.**
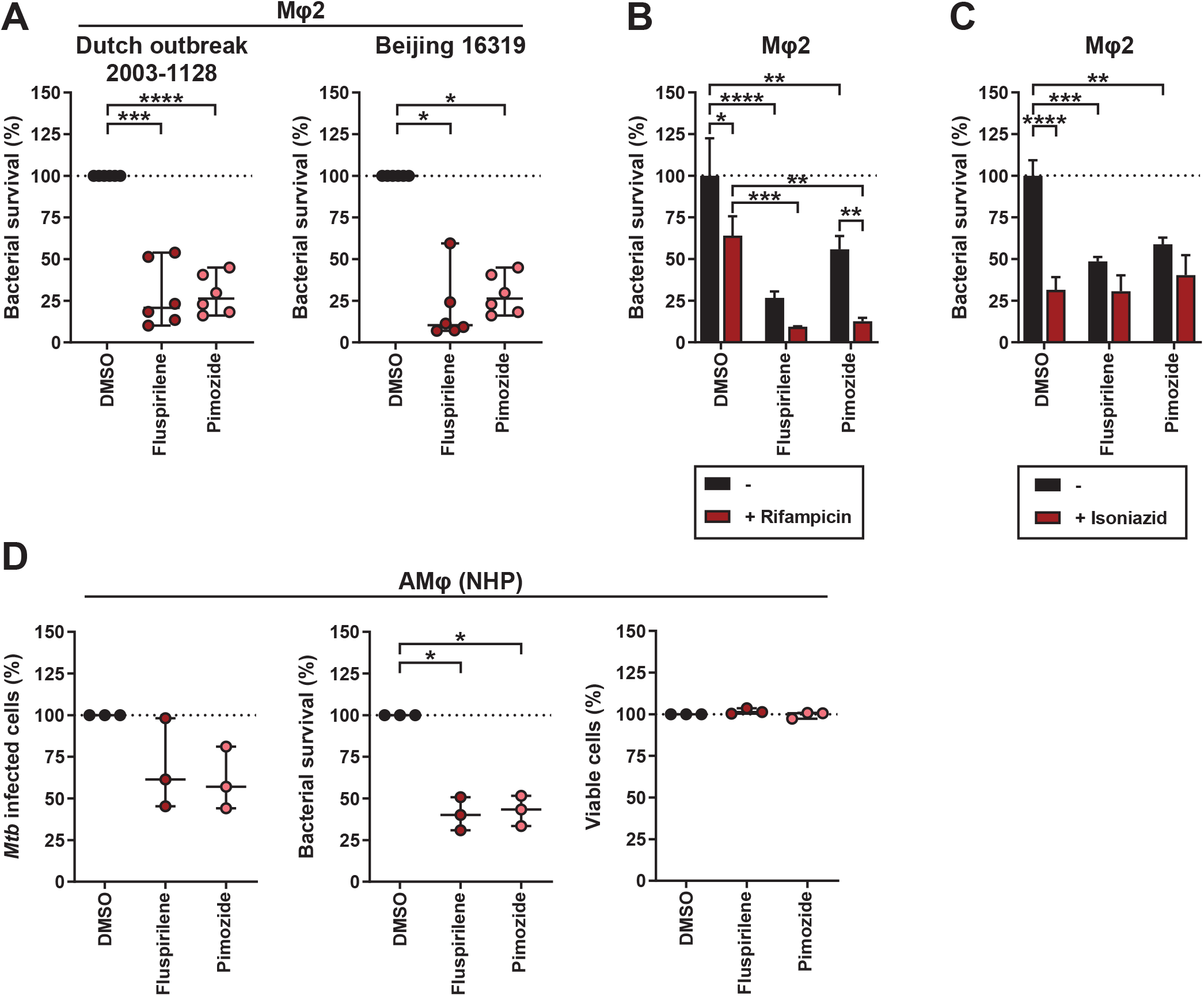
Further in depth and translational validation of Fluspirilene and Pimozide HDT activity against intracellular *Mtb*. **A.** CFU assay of Mφ2 infected with MDR-*Mtb* strain Dutch outbreak 2003-1128 (left panel) or MDR-*Mtb* strain Beijing 16319 (right panel) and treated with 10 µM of Fluspirilene, Pimozide or DMSO at equal v/v for 24 hours. Each dot represents a single donor (6 donors in total) and depicts the mean of 3 replicates. Dotted lines indicate DMSO set at 100% and median + 95% confidence intervals are shown for every condition. Statistical significance was tested using RM one-way ANOVA with Dunnett’s multiple comparison test (left panel) (*** = p-value <0.001 and **** = p-value <0.0001) or Wilcoxon matched-pairs signed rank test with post-hoc Benjamini-Hochberg correction (right panel) (* = q-value <0.1). **B.** CFU assay of Mφ2 infected with *Mtb* and treated with a combination of a suboptimal dose of Rifampicin (0.05 µg/ml) and 10 μM of Fluspirilene, Pimozide or DMSO at equal v/v for 24 hours. Bars depict mean ± standard deviation of 3 replicates from a representative donor (out of 4 donors tested), expressed as a percentage of the DMSO control in the absence of Rifampicin. Black bars represent Fluspirilene, Pimozide or DMSO only, red bars represent the combination with Rifampicin. Statistical significance was tested using a two-way ANOVA with Tukey’s multiple comparisons test comparing Fluspirilene or Pimozide treatment (in the absence or presence of Rifampicin) to the corresponding DMSO control (* = p-value <0.05, ** = p-value <0.01, *** = p-value <0.001 and **** = p-value <0.0001). **C.** CFU assay of Mφ2 infected with *Mtb* as in **B**, but a suboptimal dose of Isoniazid (0.4 µg/ml) was used. Bars depict mean ± standard deviation of 3 replicates from a representative donor (out of 3 donors tested), expressed as a percentage of the DMSO control in the absence of Isoniazid. **D.** Flow cytometry assay measuring the percentage of *Mtb*-infected cells (left panel), CFU assay (middle panel) and cell viability assay measuring the percentage of live cells (right panel) of *ex vivo* non-human primate bronchoalveolar lavage cells enriched for alveolar macrophages (NHP AMφ) infected with DsRed-expressing *Mtb* and treated with 10 µM of Fluspirilene, Pimozide or DMSO at equal v/v for 24 hours. Each dot represents a single NHP donor (3 donors in total) and depicts the mean of 1 to 2 replicates (left panel), 6 replicates (middle panel) or 3 replicates (right panel), expressed as a percentage of the DMSO control. Statistical significance was tested using a RM one-way ANOVA with Dunnett’s multiple comparison test (* = p-value <0.05).

Clinical application of HDT will most likely be considered as adjunct to standard antibiotic therapy for TB [14]. Since HDT and antibiotics have different targets, we investigated whether synergistic/additive effects between these treatments could be detected. Combining either Fluspirilene or Pimozide with a suboptimal dose of Rifampicin (0.05 µg/ml) inhibited *Mtb* outgrowth to a significantly larger extent compared to Fluspirilene, Pimozide, or Rifampicin treatment alone (**Figure 2B**). Unexpectedly, this effect was not observed when Fluspirilene and Pimozide were combined with a suboptimal dose of Isoniazid (0.4 µg/ml) (**Figure 2C**). While not further investigated here, a possible explanation for this could be that the diphenylbutylpiperidines alter host cell processes required for Isoniazid’s conversion into its active form [48].

To further strengthen the potential utility of Fluspirilene and Pimozide, we next tested their efficacy on *Mtb-*infected *ex vivo* non-human primate (NHP) bronchoalveolar lavage (BAL) cells which are a rich source of alveolar macrophages (AMφ), the primary target cells for *Mtb*. Cross-species efficacy of the drugs would suggest functional conservation of their host targets as well as translate and validate our above findings in human cells. Importantly, 24 - hour treatment with either one of the drugs decreased outgrowth of intracellular *Mtb.* This was reflected in the reduced percentage of infected cells **(Figure 2D, left panel**), and especially in the reduced number of colony forming bacteria (**Figure 2D, middle panel**). Notably, Fluspirilene and Pimozide treatment showed noy toxicity towards the *ex vivo* NHP AMφ (**Figure 2D, right panel**).

Taken together, these results significantly strengthen the potential value of Fluspirilene and Pimozide in HDT approaches against intracellular infections, including MDR-*Mtb* bacteria, and demonstrate their activity also in (NHP) infected alveolar macrophages.

### Fluspirilene and Pimozide regulate autophagy, induce a lysosomal response and enhance bacterial presence in autophago(lyso)somes

Because Fluspirilene and Pimozide are known to modulate and induce autophagy [49], we investigated this as a potential mode of action against intracellular *Mtb*. Mφ2 were infected with *Mtb*, treated for 4 hours with Fluspirilene and Pimozide, stained with CYTO-ID (a tracer that stains all autophagy-related vesicles) and visualized using confocal microscopy (**Figure 3A, left panel**). Importantly, both Fluspirilene and Pimozide, although not statistically significant, tended to increase the CYTO-ID vesicle area and the bacterial localization in these vesicles (**Figure 3A, middle and right panel**), lending support to the involvement of autophagy in Fluspirilene and Pimozide’s mode of action against intracellular *Mtb* in infected human cells.

**Figure 3.**
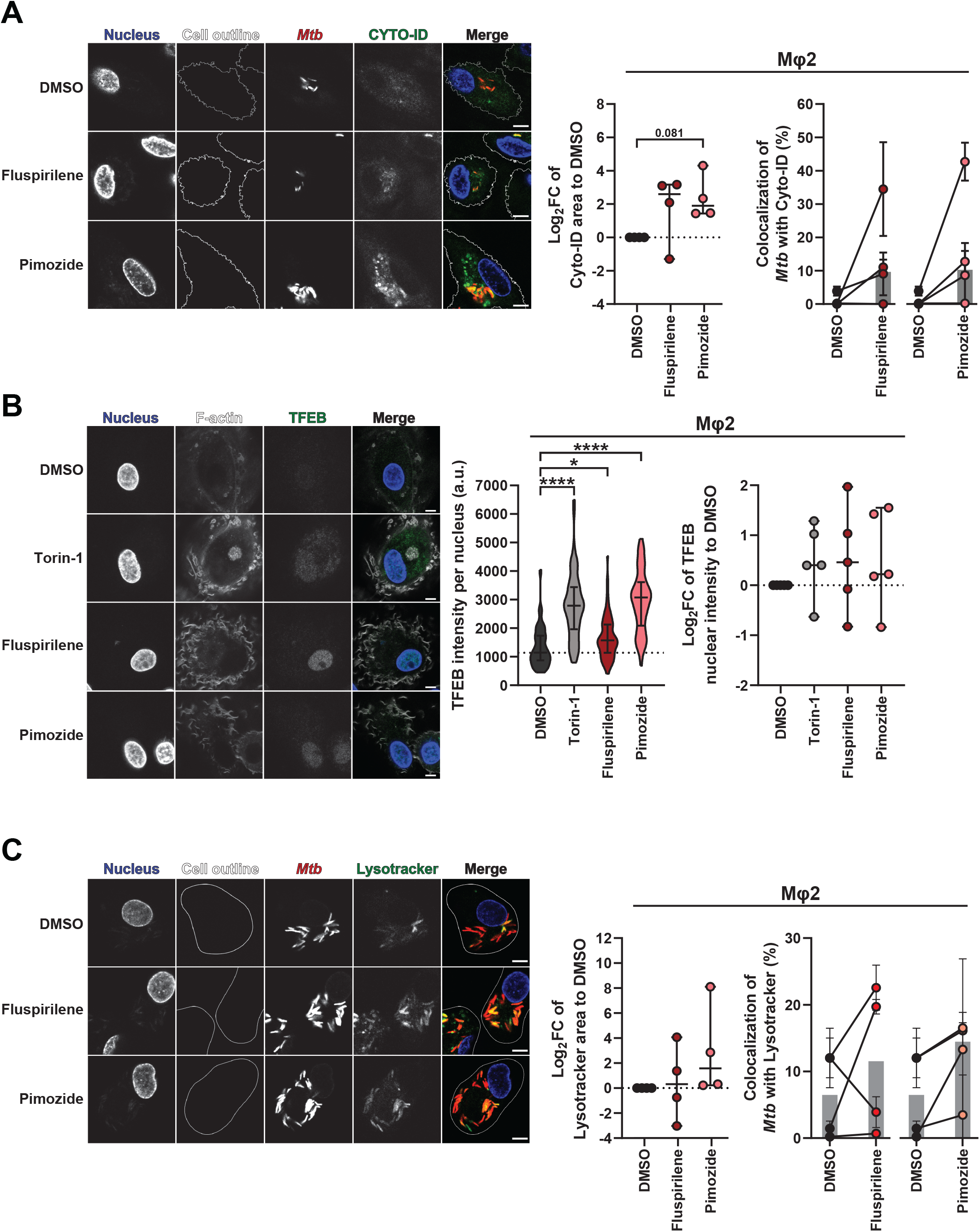
Fluspirilene and Pimozide regulate autophagy, induce a lysosomal response and enhance bacterial presence in autophago(lyso)somes. **A.** Confocal microscopy of DsRed-expressing *Mtb*-infected Mφ2 treated with 10 µM of Fluspirilene, Pimozide, 1 µM Torin-1 or DMSO at equal v/v for 4 hours. 30 min prior to the experimental endpoint cells were incubated with CYTO-ID to stain for autophagy-related vesicles, fixed and counterstained for the nucleus using Hoechst 33342. Shown in the left panel are representative images of Mφ2, while the middle panel displays the quantification of CYTO-ID positive areas (log_2_FC CYTO-ID area to DMSO) and the right panel the quantification of *Mtb* colocalization with CYTO-ID positive vesicles. Scale bar annotates 5 µm. Each dot represents a single donor (4 donors in total) and depicts the mean (middle panel) or mean ± standard deviation (right panel) of 3 replicates and with median and 95% confidence intervals shown (middle panel) or median shown by gray bars (right panel). Statistical significance was tested using a RM one-way ANOVA with Dunnett’s multiple comparison test. **B.** Confocal microscopy of DsRed-expressing *Mtb*-infected Mφ2 treated with 10 µM of Fluspirilene, Pimozide, 1 µM Torin-1 as positive control or DMSO at equal v/v for 4 hours. Cells were fixed at the experimental endpoint, permeabilized using 0.1% Triton-X, stained for TFEB and counterstained for the nucleus and F-actin using Hoechst 33342 and Phalloidin, respectively. Shown in the left panel are representative images of Mφ2, while the middle panel shows violin plots representing all nuclei quantified (n=79, n=64 n=71 and n=56 for DMSO, Torin-1, Fluspirilene and Pimozide, respectively) of a representative donor with median and interquartile range indicated and the right panel displays the log_2_FC of median nuclear TFEB intensity per donor normalized to DMSO (5 donors in total) with median and 95% confidence intervals indicated. Scale bar annotates 5 µm. Dotted line indicates median of DMSO. Statistical significance was tested using Kruskal-Wallis test with Dunn’s multiple comparison test (middle panel) (* = p-value <0.05 and **** = p-value <0.0001) or RM one-way ANOVA with Dunnett’s multiple comparison test (right panel). **C.** Confocal microscopy of DsRed-expressing *Mtb*-infected Mφ2 treated with 10 µM of Fluspirilene, Pimozide, 1 µM Torin-1 or DMSO at equal v/v for 4 hours. 30 min prior to the experimental endpoint cells were incubated with Lysotracker Deep Red to stain for acidic vesicles, fixed and counterstained for the nucleus using Hoechst 33342. Shown in the left panel are representative images of Mφ2, while the middle panel displays the quantification of Lysotracker positive areas (log_2_FC Lysotracker area to DMSO) and the right panel the quantification of *Mtb* colocalization with Lysotracker positive vesicles. Scale bar annotates 5 µm. Each dot represents a single donor (4 donors in total) and depicts the mean ± standard deviation of 3 replicates. Shown are median and 95% confidence intervals (middle panel) or median by gray bars (right panel). Statistical significance was tested using a RM one-way ANOVA with Dunnett’s multiple comparison test.

To assess whether the reduction in *Mtb* outgrowth induced by Fluspirilene and Pimozide could also be associated with an increase in lysosomal activity we first examined the nuclear accumulation of transcription factor EB (TFEB), a master regulator of the <u>c</u>oordinated lysosomal <u>e</u>xpression <u>a</u>nd regulation (CLEAR) gene network, as well as autophagy [50, 51]. Nuclear intensity of TFEB was quantified in *Mtb*-infected Mφ2 after 4 hours of treatment with Fluspirilene, Pimozide or Torin-1 as positive control [52], using confocal microscopy (**Figure 3B, left panel**). A highly significant increase in nuclear TFEB was observed for both Fluspirilene and Pimozide, even to a similar extent as the positive control Torin-1 (**Figure 3B, right panel**). To investigate the lysosomal response in more detail we employed the lysosomal tracer Lysotracker. DsRed-*Mtb*-infected Mφ2 were treated with Fluspirilene and Pimozide for 4 hours and the lysosomal area and localization of bacteria in lysosomes was quantified (**Figure 3C, left panel**). Pimozide tended to induce an increase in lysosomal area, in contrast to Fluspirilene (**Figure 3C, middle panel**). Additionally, Fluspirilene and Pimozide induced a mild, though statistically not significant average increase in colocalization of bacteria and lysosomes compared to DMSO (**Figure 3C, right panel**).

Collectively, these results suggest that the lysosomal response key regulator TFEB and likely also the autophagic response are involved in enhancing host defense induced by diphenylbutylpiperidines, which restricts outgrowth of intracellular *Mtb*. Notwithstanding, the observed limited effect sizes for some of these processes suggested to us that these mechanisms likely do not fully account for the potent HDT activity of Fluspirilene and Pimozide against intracellular *Mtb*. We therefore postulated that other mechanisms of action are likely involved in the control of intracellular bacteria upon treatment with Fluspirilene and Pimozide.

### Fluspirilene and Pimozide inhibit STAT5 function and Pimozide additionally reduces the presence of cytokine-inducible SH2-containing protein (CISH) on *Mtb* phagosomes

Because Pimozide has been reported to inhibit STAT5 function by dephosphorylation [53], and work by Queval, Song [54] recently uncovered a STAT5-mediated control of phagosomal acidification induced by *Mtb*, we explored functional inhibition of STAT5 by diphenylbutylpiperidines as a potential mode of action to control intracellular *Mtb*. First, the effect of 4-hour treatment with diphenylbutylpiperidines on nuclear presence of phosphorylated STAT5 (P-STAT5) was investigated in *Mtb*-infected Mφ2 using confocal microscopy (**Figure 4A, left panel**). Both Fluspirilene and Pimozide significantly decreased the nuclear presence of P-STAT5 **Figure 4A, middle and right panel**). To further corroborate our finding that Fluspirilene and Pimozide lowered nuclear P-STAT5 quantity and hence its transcriptional activity, we analyzed the expression levels of cytokine-inducible SH2-containing protein (*CISH*), a transcriptional target of STAT5 [55]. We confirmed the observation by Queval, Song [25] that *Mtb* infection increases *CISH* transcript levels in our *Mtb*-Mφ2 model after 4 hours (**Figure 4B**). Unexpectedly, Fluspirilene increased *CISH* transcript levels, even to a higher extent than the positive control GM-CSF [56], while Pimozide did not affect *CISH* transcript expression levels, despite the compound’s’ effects on P-STAT5 (**Figure 4B)**. We therefore quantified colocalization of CISH protein with *Mtb*, since the presence of CISH on the *Mtb* phagosome has been associated with bacterial survival [54]. Pimozide treatment caused a significant decrease of CISH fluorescence intensity per bacterium within the 4h time-period, but Fluspirilene did not clearly show a difference compared to DMSO (**Figure 4C**).

**Figure 4.**
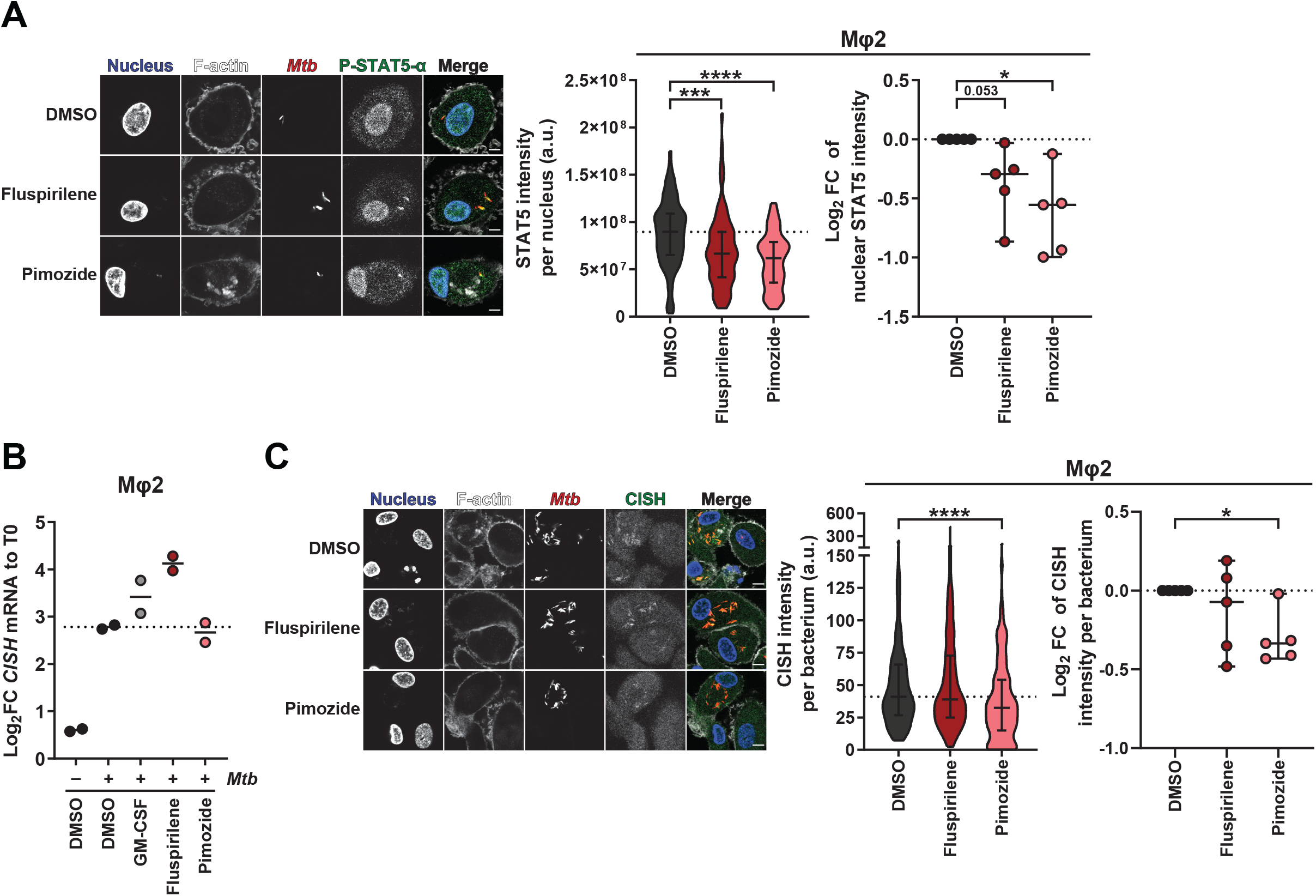
Fluspirilene and Pimozide inhibit STAT5 function and Pimozide additionally reduces the presence of cytokine-inducible SH2-containing protein (CISH) on *Mtb* phagosomes A. Confocal microscopy of DsRed-expressing *Mtb*-infected Mφ2 treated with 10 µM of Fluspirilene, Pimozide or DMSO at equal v/v for 4 hours. Cells were fixed at the experimental endpoint, permeabilized using 0.1% Triton-X, stained for P-STAT5 and counterstained for the nucleus and F-actin using Hoechst 33342 and Phalloidin, respectively. Shown in the left panel are representative images of Mφ2, while the middle panel shows violin plots representing all nuclei quantified (n=73, n=77 and n=75 for DMSO, Fluspirilene and Pimozide, respectively) of a representative donor with median and interquartile range indicated and the right panel displays the log_2_FC of median nuclear P-STAT5 intensity per donor normalized to DMSO (5 donors in total) with median and 95% confidence intervals indicated. Scale bar annotates 5 µm. Dotted line indicates median of DMSO. Statistical significance was tested using Kruskal-Wallis test with Dunn’s multiple comparison test (middle panel) (** = p-value <0.01, *** = p-value <0.001 and **** = p-value <0.0001) or RM one-way ANOVA with Dunnett’s multiple comparison test (* = p-value <0.05). **B.** Mφ2 derived from 2 donors were mock- or *Mtb*-infected and *Mtb*-infected Mφ2 were subsequently treated with 10 µM of Fluspirilene, Pimozide, GM-CSF (50 ng/ml) as positive control or DMSO at equal v/v for 4 hours. Transcript levels of Cytokine-inducible SH2-containing protein (CISH) were determined in duplicate using qRT-PCR before (0h baseline) and 4-hour post-infection. Data was normalized to GAPDH (ΔCt). Each dot represents a single donor and displays the log_2_FC expression levels of CISH in response to treatment compared to their respective baseline control (T0) (ΔΔCt). Horizontal lines indicate median expression levels and dotted line indicates the median of *Mtb*-infected Mφ2 treated with DMSO. **C.** Confocal microscopy of DsRed-expressing *Mtb*-infected Mφ2 treated with 10 µM of Fluspirilene, Pimozide or DMSO at equal v/v for 4 hours. Cells were fixed at the experimental endpoint, permeabilized using 0.1% Triton-X, stained for CISH and counterstained for the nucleus and F-actin using Hoechst 33342 and Phalloidin, respectively. Shown in the left panel are representative images of Mφ2, while the middle panel shows violin plots representing all bacteria quantified (n=182, n=179 and n=217 for DMSO, Fluspirilene and Pimozide, respectively) of a representative donor with median and interquartile range indicated and the right panel displays the log_2_FC of median CISH integrated intensity per bacterium per donor normalized to DMSO (5 donors in total) with median and 95% confidence intervals indicated. Scale bar annotates 5 µm. Dotted line indicates median of DMSO. Statistical significance was tested using Kruskal-Wallis test with Dunn’s multiple comparison test (middle panel) (**** = p-value <0.0001) or RM one-way ANOVA with Dunnett’s multiple comparison test (* = p-value <0.05).

Taken together, these data suggest that Fluspirilene and Pimozide both decrease P-STAT5 protein levels in the nucleus. In addition, although the effects on intracellular *Mtb* bacteria did not correlate with differential expression of the STAT5 target *CISH* at the transcript level, Pimozide lowered CISH protein presence on the *Mtb* phagosome. Thus, both Fluspirilene and Pimozide likely target intracellular *Mtb* by a mechanism involving P-STAT5, which for Pimozide, but not Fluspirilene, includes routing through the CISH effector pathway. These results suggest that structurally highly related HDT compounds can nevertheless subtly differ in their mode of actions against intracellular infection.

### Pimozide induces ROS/RNS production and antioxidants impair bacterial killing by Pimozide as well as Fluspirilene

Next to autophagy and lysosomal degradation, reactive oxygen species (ROS) and reactive nitrogen species (RNS) represent additional mechanisms that could play a key role in controlling intracellular *Mtb*. Of note, Pimozide has been reported to induce ROS production in various cell types [57–59], prompting us to explore the role of radical species in the mode of action of both Fluspirilene and Pimozide. Using a general oxidative stress indicator (CM-H2DCFDA), we measured ROS/RNS production in uninfected Mφ2 (due to biosafety level 3-restrictions) during treatment with Fluspirilene and Pimozide. Pimozide but not Fluspirilene treatment clearly increased ROS/RNS production (**Figure 5A and 5B, left panel**). To confirm that probe conversion was caused by cellular produced radical species, the probe was incubated with Pimozide in a cell-free system, which showed no significant effect (**Figure 5C**). To examine whether we could confirm the induction of ROS/RNS production by Pimozide, we added the antioxidant N-acetylcysteine (NAC) to the Pimozide treatment regimen (**Figure 5B, right panel**). In 7 out of 9 donors Pimozide-induced ROS/RNS production was impaired by the addition of NAC, indicating that the probe conversion indeed reflects elevated ROS/RNS production.

**Figure 5.**
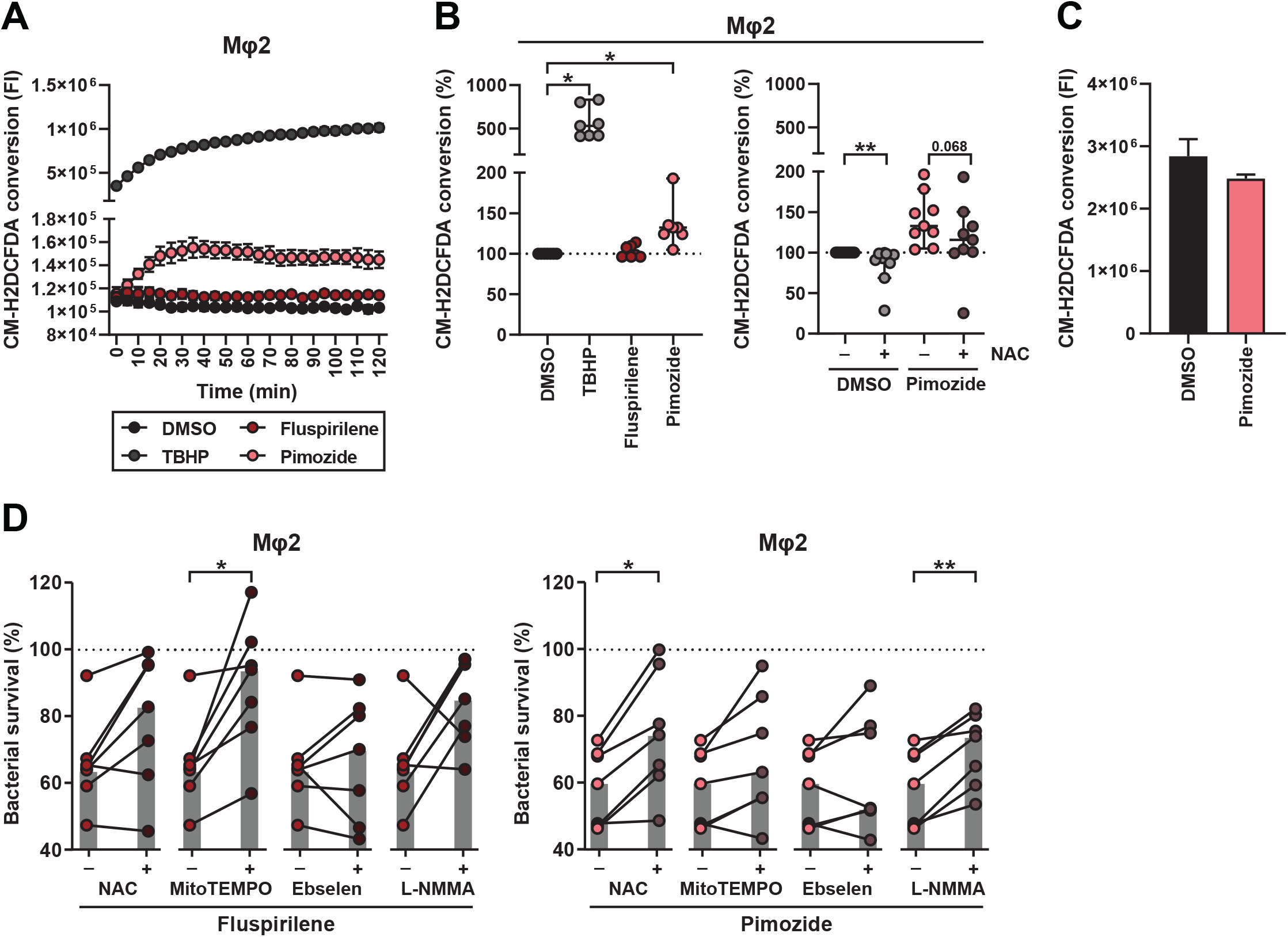
Pimozide induces ROS/RNS production and antioxidants impair bacterial killing by Pimozide as well as Fluspirilene. **A.** Mφ2 were pulsed for 30 min with 5 µM of probe CM-H2DCFDA followed by exposure to 10 µM Fluspirilene or Pimozide, 50 µM TBHP as positive control, or DMSO at equal v/v. Production of reactive oxygen species (ROS) was monitored by measuring Fluorescence intensity (522 nm) over a time course of two hours. Each dot depicts the mean of 3 replicates of a representative donor (7 donors in total). **B.** Mφ2 were pulsed for 30 min with 5 µM of probe CM-H2DCFDA followed by exposure to 10 µM Fluspirilene or Pimozide, 50 µM TBHP as positive control or DMSO at equal v/v for the duration of the experiment (left panel). Pimozide and DMSO were additionally combined with 5 mM of the antioxidant N-acetyl cysteine (NAC) (right panel). Each dot represents the area under the curve (AUC) of a single donor (7 donors in left panel and 9 donors in right panel) and depicts the mean of 3 to 6 replicates. Median with 95% confidence intervals are shown. Statistical significance was tested using a Wilcoxon matched-pairs signed rank test with post-hoc Benjamini-Hochberg correction (left panel) (* = q-value <0.1) or Wilcoxon matched-pairs signed rank test (right panel) (** = p-value <0.01). **C.** Probe CM-H2DCFDA (5 µM) fluorescence measured in the presence of 10 µM of Pimozide or DMSO at equal v/v in the absence of cells. Bars display the mean ± standard deviation of 3 replicates and represent the fluorescence intensity measured after 60 min of incubation in one single experiment. Statistical significance was tested using an unpaired t test with Welch’s correction. **D.** CFU assay of *Mtb*-infected Mφ2 and treated with 10 µM of Fluspirilene (left panel) or Pimozide (right panel) combined with anti-oxidants (5 mM of NAC, 10 µM of MitoTempo, 25 µM of Ebselen or 1 mM of L-NMMA) or DMSO at equal v/v for 24 hours. Each dot represents a single donor (7 donors in total) and depicts the mean of 3 to 6 replicates. Dotted lines indicate DMSO set at 100% with median indicated by gray bars. Statistical significance was tested using a RM one-way ANOVA with Dunnett’s multiple comparison test (* = p-value <0.05).

To study the importance of Pimozide-induced ROS/RNS production in its control of intracellular bacterial infection, *Mtb*-infected Mφ2 were treated with either Pimozide or Fluspirilene in combination with various ROS/RNS inhibitors, namely: NAC, Ebselen (a NOX inhibitor and glutathione peroxidase mimic [60]), MitoTEMPO (a mitochondria-targeted radical scavenger [61]), and L-NMMA (a nitric oxide synthase (NOS) inhibitor [62]) (**Figure 5D**). NAC, and L-NMMA, but not Ebselen and MitoTEMPO, significantly inhibited the HDT effects of Pimozide (**Figure 6D, right panel**). Interestingly, we also observed a significant effect of MitoTEMPO but not NAC, L-NMMA or Ebselen, on Fluspirilene (**Figure 5D, left panel**), indicating a -possibly differential- role for ROS/RNS in the mode of action of both Fluspirilene and Pimozide. NAC, MitoTEMPO, L-NMMA and Ebselen treated *Mtb*-infected Mφ2 all showed some reduced bacterial outgrowth, which was significant for NAC and Ebselen (**Supplementary Figure 1**), further emphasizing the role of ROS/RNS in intracellular bacterial control, which is supported by their antagonistic effect on Pimozide and Fluspirilene induced infection control.

**Figure 6.**
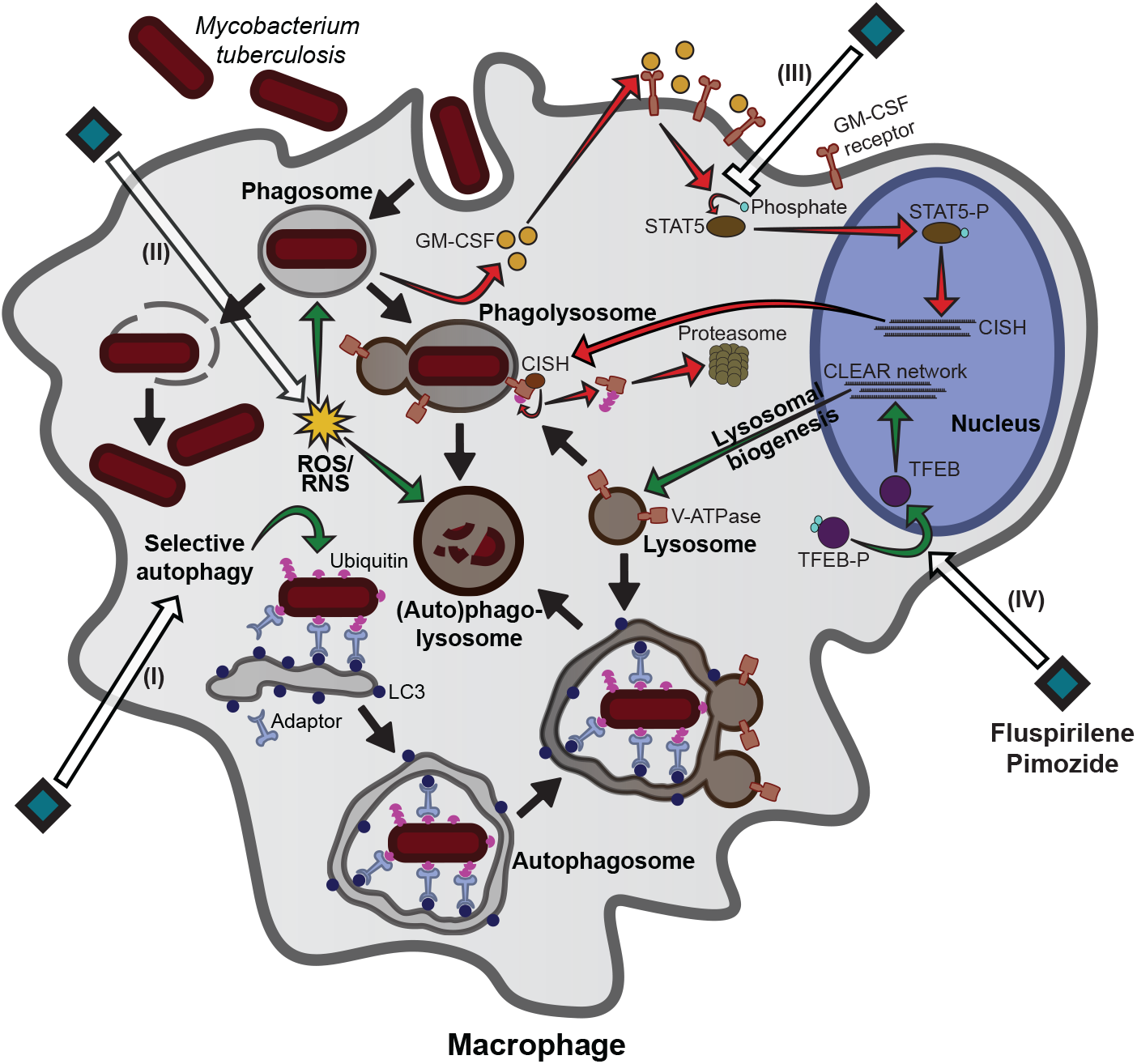
Model of the modes of action of Fluspirilene and Pimozide. Both Fluspirilene and Pimozide increased the localization of *Mtb* in autophagy-related vesicles implying they induce selective autophagy, i.e. xenophagy (I). Although increased ROS/RNS production was significantly induced by Pimozide only, antioxidants impaired the efficacy of Pimozide and to a lesser extent also Fluspirilene (II). Fluspirilene and Pimozide both inhibited STAT5 activity and Pimozide consequently reduced CISH localization on the *Mtb* containing vesicle (III). Lastly, Fluspirilene and to a larger extent Pimozide, increased nuclear TFEB localization concomitant with an increased lysosomal response (IV). Black arrows indicate the general process of *Mtb* phagocytosis and subsequent events, white arrows show compound mediated activation or inhibition of targets and green and red lines depict signaling pathways and protein interactions with reduced or increased bacterial outgrowth as outcome, respectively.

In conclusion, we have identified two clinically approved drugs, Fluspirilene and Pimozide, which we show to have novel and significant potential as host-directed therapeutics for the treatment of intracellular bacterial pathogens such as *Mtb* and *Stm*, mechanistically operating by multiple mechanisms of action. We show that both Fluspirilene and Pimozide significantly alter the lysosomal response and tend to increase autophagic targeting of *Mtb*, likely via nuclear translocation of TFEB. Furthermore, our findings support two additional modes of action, particularly associated with Pimozide-mediated activity. Firstly, Fluspirilene and Pimozide counteract *Mtb*-induced STAT5 phosphorylation, but only Pimozide thereby reduced the presence of CISH on the *Mtb* phagosome, thus probably regulating its acidification [54]. Secondly, Pimozide induced measurable ROS/RNS production and its efficacy was impacted by ROS/RNS inhibitors. Together these mechanisms likely work in concert (**Figure 6**) to reduce intracellular bacterial outgrowth and thus could be targeted by adjunct HDT in difficult to treat intracellular infections such as *Mtb* and *Stm* and likely also related pathogens.

## Discussion

In this study, we aimed to find new host-directed therapy (HDT) candidate drugs for the treatment of (drug resistant) TB and other difficult to treat intracellular bacterial infections, and identified two structural analogues of the diphenylbutylpiperidine-class, Fluspirilene and Pimozide, with strong efficacy against intracellular *Mtb* and *Stm*. We next validated their HDT potential in *Mtb* and *Stm* primary human macrophages with either pro- or anti-inflammatory phenotypes. Importantly, we corroborated their activity against intracellular *Mtb* in a cross-species non-human model using bronchoalveolar lavage fluid macrophages, which are a rich source of alveolar macrophages (AMφ), the primary target cells for *Mtb*. This cross-species efficacy underscored functional conservation of host target molecules and independently validated their effects in human *Mtb* infected cells. Additionally, the strong HDT activity of Fluspirilene and Pimozide was confirmed also against clinical isolates of multi-drug resistant (MDR)-*Mtb* strains, and moreover acted in a synergistic and/or additive mode with a suboptimal dose of Rifampicin. Extracellular bacterial proliferation itself was unaffected by these compounds, indicating that their targeting of host mechanisms is responsible for their activity. Furthermore, the high efficacy shown against *Stm*, indicates that these drugs could be repurposed as treatment against a wider range of intracellular pathogens as already shown for other HDTs and pathogens [63].

Interestingly, we found several modes of action employed by Fluspirilene and Pimozide, including modulation of the lysosomal response, autophagy, induction of ROS/RNS production and finally the inhibition of STAT5 with consequently decreased localization of CISH protein on the *Mtb* phagosome. We postulate that these mechanisms cooperate in concert via multiple pathways to reduce intracellular bacterial outgrowth. Drugs acting via multiple effector mechanisms might be desirable over single mechanism of action (MOA), because developing bacterial resistance against multiple host effector mechanisms simultaneously will be extremely hard to achieve (summarized in **Figure 6**).

Fluspirilene and Pimozide are potent inhibitors of several dopamine receptors and calcium channels and are used clinically to treat psychotic disorders, like schizophrenia, psychosis and Tourette syndrome [41, 64]. Some antipsychotic drugs have already been shown to exert (host directed) antimicrobial activity. For example, phenothiazine-derived drugs, which include Trifluorperazine, as well as butyrophenone derivatives, with Haloperidol as a well-known example, have therapeutic potential against several pathogens as we and others have shown previously [22, 26, 65, 66]. The diphenylbutylpiperidine-class of drugs has also been studied, revealing host-directed effects of Pimozide against both facultative and obligate intracellular pathogens, *Listeria monocytogenes* and *Toxoplasma gondii* respectively [67, 68]. Also, Fluspirilene has been shown to potentiate antimicrobial activity against several bacteria, but only in a direct antibiotic manner [69, 70]. Despite these observations, we found no direct antimicrobial effect of Fluspirilene (or Pimozide) on either *Mtb* or *Stm*. This is likely reconciled by the fact that Fluspirilene displayed minimum inhibitory concentrations (MICs) of > 80 µM for *E. coli* and *K. pneumoniae* which both belong to the same order as *Stm*, namely Enterobacterales, whereas in our work we employed significantly lower (10 µM) concentrations at which strong HDT but no direct anti-bacterial effects were observed [69].

Our work suggests that four different host effector mechanisms are likely to play a role in the treatment-efficacy of Fluspirilene and Pimozide. A first mechanism we demonstrated was that Fluspirilene and Pimozide activated autophagy in *Mtb*-infected Mφ and increased the localization of *Mtb* in autophagic vesicles (**Figure 3A**). Although this was not investigated in further detail, we hypothesize that Ca^2+^ signaling could be important to this effector mechanism. Both Fluspirilene and Pimozide antagonize calcium channel activity and can lower intracellular Ca^2+^ levels, thereby inactivating Calpain that leads to the induction of autophagy [41, 64, 71, 72]. In support of this, a screen of Fluspirilene analogues showed that Ca^2+^ channel inhibition was necessary for their ability to induce autophagy in a neuroglioma cell line, since analogues that lacked the ability to block Ca^2+^ channels lacked autophagy-inducing activity [71]. Importantly, the host Ca^2+^ signaling pathway is also manipulated directly by *Mtb* to inhibit autophagy [73]. Whether Ca^2+^-related effects also play a role in HDT activity of Fluspirilene and Pimozide against intracellular *Mtb* and *Stm,* however, remains currently unknown and will require future studies. Recently, new diphenylbutylpiperidine analogues with stronger autophagy-inducing capabilities were developed that represent interesting novel HDT for future studies [74].

The second mechanism that Fluspirilene and Pimozide are likely to employ is the lysosomal response (**Figure 3**). An additional new finding with mechanistic implications was that treatment with Fluspirilene and Pimozide strongly increased the intranuclear presence of TFEB (**Figure 3B**). This transcriptional regulator controls a coordinated lysosomal response [75], as well as autophagy [76]. This ability of Fluspirilene and Pimozide was, to the best of our knowledge, hitherto unknown. TFEB translocation has been observed for drugs of the phenothiazine-class including Trifluorperazine [77], suggesting this might represent common functionality of several different antipsychotic drugs.

The third mechanism we uncovered was phagosomal acidification, a process vital in the host response against intracellular bacteria, which is inhibited by the protein CISH that is under transcriptional control of STAT5 [54]. This pathway is exploited by *Mtb* which induces GM-CSF to activate STAT5, leading to enhanced *CISH* expression and consequent V-ATPase degradation [54]. Although treatment with both Fluspirilene and Pimozide led to a reduction of nuclear STAT5, we could experimentally confirm the decrease in CISH protein on *Mtb* containing vesicles only for Pimozide (**Figure 4A and C**). Further research will be needed to further unravel the subtle differences in precise MOAs between Pimozide and Fluspirilene, but this is beyond the scope of the current work.

As a fourth effector mechanism, we found that Pimozide strongly induced ROS/RNS production. Mechanistically, the efficacy of both Pimozide and Fluspirilene towards intracellular *Mtb* was inhibited by ROS/RNS inhibitors. Of these two diphenylbutylpiperidines, only Pimozide is known to induce ROS/RNS production and to inhibit the expression of antioxidant genes, like catalase, *in vitro* as well as *in vivo* [57, 78]. Interestingly, similar to the results from Cai, Zhou [57], addition of n-acetyl-cysteine (NAC), a general ROS scavenger, partly inhibited the Pimozide induced increase in ROS/RNS levels by human macrophages, and reduced bacterial survival (**Figure 5B and D**). Since multiple signaling pathways could contribute to the ROS/RNS production caused by Pimozide, we investigated this in more detail. The respiratory burst induced upon phagocytosis did not seem to be involved since Ebselen, a NADPH oxidase (NOX) inhibitor [79], did not restore bacterial survival. Because L-NMMA, selective for nitric oxide synthase (NOS) affected Pimozide’s efficacy, a role for nitric oxide (NO) seems likely. It has become evident that the functions of ROS and NO are intertwined and that inhibition of one leads to cross-inhibition of the other, supporting this possibility [80, 81].

Taken together, we have identified four potential mechanistic effector pathways each of which is regulated by Fluspirilene and/or Pimozide, and which likely act in concert to reduce intracellular bacterial outgrowth. Combinatorial treatment regimens using multiple approaches including HDT as well as anti-bacterial molecules are considered indispensable to eradicating TB [82]. An interesting finding was that although Fluspirilene and Pimozide displayed additive effects with a low dose regimen of Rifampicin, this was not observed with Isoniazid. Isoniazid is a prodrug and requires conversion by the *Mtb* catalase-peroxidase (KatG) enzyme to its active form, resulting in the production of bacteria-derived reactive species [48, 83]. These radicals have both direct and indirect antimicrobial properties and in the latter case host ROS/RNS production can lead to the induction of autophagy, which is capable of targeting *Mtb* [48].

Since the continuing rise of global drug resistance will affect the treatment of many bacterial infections, approaches complementary to bacterial-directed strategies can offer important supplemental approaches. We have demonstrated the novel HDT potential of two approved antipsychotic drugs, Fluspirilene and Pimozide, against *Mtb* and *Stm,* two important but unrelated intracellular pathogens. Our results provide new insights into the molecular and functional effects of these two diphenylbutylpiperidines on key mechanisms of host defense, particularly: autophagy, lysosomal acidification, ROS/RNS generation and phagosome maturation via STAT5/CISH inhibition. We show that Fluspirilene and Pimozide target these host responses, by similar but subtly different MOAs, which is accompanied by reduced intracellular outgrowth of *Mtb* and *Stm* in human cells. Based on these findings, we propose that the class of diphenylbutylpiperidines could be repurposed as novel HDT drugs for both TB and salmonellosis and further explored as host-directed compounds against other intracellular pathogens.

## Supplementary Figure Legends

**Supplementary Figure 1.**
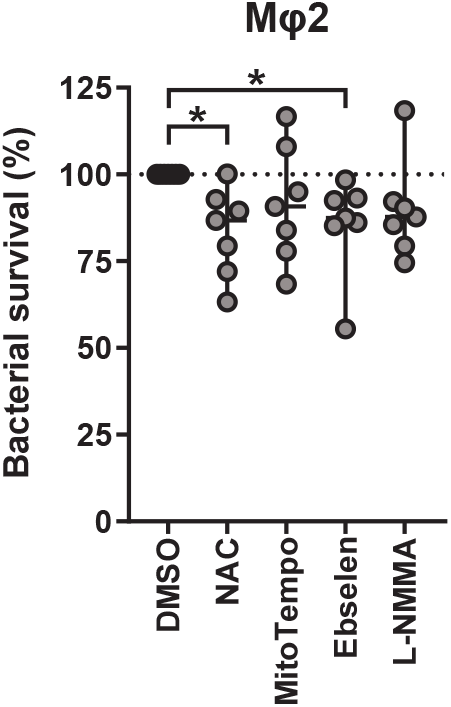
Effect of ROS/RNS inhibitors on *Mtb* outgrowth. CFU assay of *Mtb*-infected Mφ2 and treated with antioxidants (5 mM of NAC, 10 µM of MitoTempo, 25 µM of Ebselen or 1 mM of L-NMMA) or DMSO at equal v/v for 24 hours. Each dot represents a single donor (7 donors in total, as in Figure 6D) and depicts the mean of 3 to 6 replicates. Dotted lines indicate DMSO set at 100% with median indicated by gray bars. Statistical significance was tested using Wilcoxon matched-pairs signed rank test with post-hoc Benjamini-Hochberg correction (* = q-value <0.1).

## Acknowledgements

We gratefully acknowledge Dr J. Bestebroer (VUMC, Amsterdam, The Netherlands) for mycobacterial reporter constructs and Dick van Soolingen and Kirsten Kremer (RIVM, Bilthoven, The Netherlands) for providing the MDR-*Mtb* strains.

This project was funded by the European Union’s Seventh Programme for research, technological development and demonstration under grant agreement N° PhagoSys HEALTH-F4-2008-223451, grants from the Netherlands Organization for Health Research and Development (ZonMw-TOP grant 91214038) and NWO Domain Applied and Engineering Sciences (NWO-TTW grant 13259). We acknowledge the support from FAPESP (grant: 2017/03332-5) to CS fellowship. The funders had no role in study design, data collection and analysis, decision to publish, or preparation of the manuscript.

## Supplementary materials

Table S1: Results of Screen-Well Autophagy Library screen on *Mtb* infected MelJuSo cells and Table S2: Raw data of confocal microscopy experiments.

## Author contribution

MTH, CJK, JE, THMO and MCH designed the experiments. MTH, CJK, JE, CS, IG, FV, SV, KVW, CGE, KD, LW performed the experiments. MTH, CJK and SV processed the experimental data and MTH, CJK, THMO and MCH contributed to the interpretation of the results. MTH, THMO and MCH wrote the manuscript and designed the figures. FAWV contributed to the interpretation of the results and reviewed the manuscript. THMO and MCH supervised the project. All authors approved the final version of the manuscript.

